# Integrating Math Modeling, Coding, and Biology in a CURE Lab

**DOI:** 10.1101/2022.01.20.477155

**Authors:** Renee Dale, Stewart Craig

## Abstract

The development of mathematical and quantitative skills is increasingly critical for biology students. Literacy in coding, statistics, and mathematical modeling enables students to engage in systems-thinking and the analysis of large data sets. Here we present a flexible tool illustrating concepts in mathematical modeling, coding, and biology for integration into both traditional ‘cookbook’ and inquiry-driven labs for freshmen biology students. We developed simple and complex mathematical models of potato catechol oxidase, a popular system to teach enzyme kinetics in undergraduate biology labs. We integrated both models into a freely-available web app for simulation and parameter estimation. The models are usable even if experimental details are unknown or poorly controlled, so that even novice students can work through the problem. We illustrate this by estimating the kinetic parameters of catechol oxidase in the complex model using data obtained from two sections of a course-based undergraduate research experience (CURE) freshman biology lab. Worksheets with questions motivating model building and simulation comprehension are provided. The effect of these exercises on students’ opinions of math, biology, and coding are evaluated using pre- and post-test surveys and student feedback. Our results show that these tools illustrate enzyme kinetics mathematically, students are not intimidated by the degree of math or coding involved, and in some cases are interested to do more, despite being unaware of the focus of the lab when signing up.

## 1 Introduction

Despite the increasing ubiquity of mathematics and quantitative skills in the life sciences, many undergraduate students matriculate without familiarity in their application. The ability to understand mathematical models is increasingly useful in academic and industrial sciences, from public health to basic research. Mathematical modeling and computational approaches help build the skills of computational and systems thinking, while simulators allow inquiry-driven science and hypothesis formation while reducing time required for experimentation [1, **?**, 2].

The importance of improving the quantitative coverage of undergraduate biology education and interdisciplinary work is paramount [5, 4]. Illustrating quantitative concepts in an interdisciplinary way has been shown to complement the quantitative courses students are already taking, as well as facilitating the synthesis of complex concepts for students who may not be intrinsically motivated to learn quantitative techniques [5, 2, 6]. However, incorporating quantitative skills into existing courses remains challenging. The possible negative impact of math anxiety on students is a concern, partially due to social perceptions of mathematics and to belief that new methods are inherently more difficult [7, 8]. We addressed this by incorporating critique of math model as part of the modeling process. This approach has been recently applied with success [9].

Potato catechol oxidase is a popular laboratory exercise for student learning about enzymes, and we developed our quantitative materials around the existing experimental designs [10, 11, 12, 13]. The catechol oxidase is responsible for the oxidation of catechol, causing browning in many fruits and vegetables. Since benzoquinone (the product) is brown, the reaction progress can be tracked with a spectrophotometer. Students tracked reaction progress and modified enzyme and substrate concentrations. They then used the simulator of the simple model of catechol oxidase kinetics to obtain rough estimates of the reaction rates. After working through the derivation of the simple model, they critiqued it and considered ways it could be improved. They obtained estimates of the reaction rate by working through the R code and plotting their data with the simple model predictions. As a result of the course, we found modest improvements in students’ opinions of math and biology, and some students expressed interest in quantitative topics. After the course, we developed a more complex mathematical model, obtaining parameter estimates with student experimental data. We found that reasonable estimates could be found despite irregularities in the data, and developed a simulator of the complex model. The resulting tools should be helpful for others working to integrate quantitative methods into biology labs.

## 2 Methods

### 2.1 Study Population

The study population was a group of 35 students across two sections of an introductory biology laboratory for science majors at Louisiana State University. These sections were CURE (course-based undergraduate research experience) labs, designed by the first author to introduce students to math modeling and coding. The students had little to no prior experience with programming, and a handful had previously encountered ordinary differential equations (ODEs). The students’ majors were in biology or related fields (e.g., agriculture, natural resources). All students had little or no prior experience with mathematical models. We received Institutional Review Board (IRB) confirmation that this study did not require approval.

### 2.2 Experimental methods

An existing catechol oxidase experimental design was used [13]. Catechol oxidase is extracted from potatoes by blending skinned potatoes in a blender, straining through coffee filter and cheesecloth, and placing the extract on ice. 100 grams of potato are blended with 500 mL of filtered water. 1 mL of extract is combined with 5 mL of water and 3 mL of catechol into a 9 mL test tube and placed inside a Spectronic 20 spectrophotometer. The absorbance was tracked using spectrophotometers every 30 seconds for 180 seconds at 510 nm, after previously blanking the spectrophotometer with 8 mL water and 1 mL extract. The data used in this paper was collected over 25 student groups, across two sections and two experiment sessions. The data are provided at https://doi.org/10.6084/m9.figshare.15079383.

### 2.3 Modeling Methods

A simple differential equation model was created, considering only substrate, enzyme, and product (Eqn 5). Differential equations were then written for each of the variables in the reaction of the enzyme and catechol using non steady state mass action kinetics. This produced 7 equations with 11 parameters (Eqn 3.1). Differential equations were solved with ode in R 3.6.3 for the simulators, and ode15s with the global optimization toolbox in Matlab 2020a for parameter estimation [14]. Figure colors were chosen to be maximally distinct [15]. Code is provided as Supplemental. Two versions of a simulator were created and shared as an R Shiny web app (https://rdale1.shinyapps.io/wischubiol2018/).

### 2.4 Surveys

The study included 2 assessments. At the beginning and end of the semester, a survey containing 21 items from previously published math and biology attitudes surveys (10 questions from each plus a control question) was provided to the students as part of normal in-class work [16, 17]. Students were provided with an “opt-in” consent form giving their consent for their data to be analyzed as part of this study (one student didn’t opt-in). The survey tracked the changes in their attitudes after completing the course. The survey took around 10 minutes to complete, and the surveys are Likert-scale response surveys. Secondly, an anonymous open-ended opinion questionnaire was provided to the students at the end of the semester using an online Google Form. The purpose of the survey was to obtain information such as what they liked most or least about the course content. The data are provided at 10.6084/m9.figshare.6169523.

### 2.5 Statistical analysis

The Likert-scale pre and post test survey produces ordinal data that may not be normally distributed [18]. We checked for reasonable symmetry of pre and post test response differences using histograms (Supplemental Fig. 1). We then used Wilcoxon signed-rank test to determine significant shifts in responses using wilcox test from package rstatix (version 0.6.0) in R 3.3.6. Using a significance level of 0.05, we found 3 questions were significantly different, and a 4th question significant at the 0.06 level. We measured reliability using Chronbach’s alpha using package psych (version 2.0.12). The reliability of all survey items was 0.37, and 0.54 when adjusted using Spearman Brown, meaning moderate reliability for the 20-item test. The R code used for these analyses is provided as a Supplemental, and all package versions are annotated.

## 3 Thinking about enzymes from a modeling perspective

Enzymes interact with substrates and catalyze their reactions which produce some product.

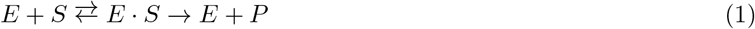

Here *E* stands for enzyme, *S* for substrate, *E·S* for the enzyme-substrate complex, and *P* for product. In this model, enzyme and substrate reversibly bind to form the enzyme-substrate complex before the enzyme catalytically releases the product.

To think about this system from a modeling perspective, students were asked to first consider what we measure during enzymatic reactions (see Supplemental worksheets). With some prompting, students may arrive at the phrase, ‘change in product over time’. Mathematically, this would be phrased as:

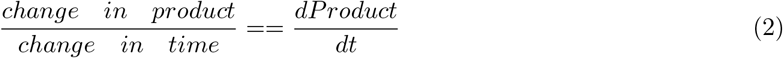

This describes the system in terms of ordinary differential equations (ODEs), or changes over time. ODEs lend themselves well to the study of biological phenomena as many biological processes are described by rates. Applying this logic to the enzyme reaction, students must identify how each of the states (enzyme, substrate, and product) are changing over time. For example, consider how the amount of substrate changes.

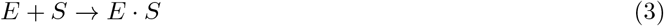

What is the directionality of this change? Does it all change immediately, or is there a rate of change (“r”)? Finally, what happens if there is no enzyme (E)? Will the substrate continue changing? Answers to these questions develop the following model, in mathematical terms:

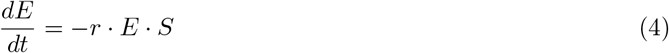

When considering biological systems from a mathematical perspective, some features need to be addressed as they may not be obvious to students. For example, enzymatic catalysis of substrates depends on the amount of substrate *and* the amount of enzyme, and the amount of substrate decreases at some *rate*. Applying these methods to the remaining states in the system students derive the following simple system of equations describing this system. The system of equations is simplified by specifying that the amount of free enzyme does not change over time. This is accurate when the amount of substrate is much larger than the amount of enzyme, and the amount of substrate isn’t exhausted (i.e., Michaelis-Menten kinetics).

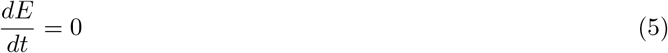

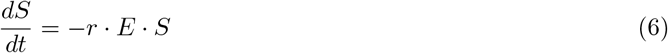

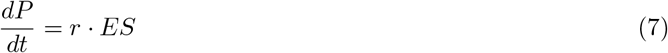

This exercise allows us to illustrate how helpful mathematical models can be, to determine if your current understanding of the system is sufficient to explain the data you see. It is important to emphasize that “all models are wrong, but some are useful” [19]. To facilitate engagement with mathematical models, a web app of this simple model was created (available at https://rdale1.shinyapps.io/wischubiol2018/). The simulator allows students to alter the amount of enzyme, substrate, or the reaction rate on the progress of the reaction. It plots the resulting changes in substrate (catechol), and product (benzoquinone) over time. Students were asked to modify the reaction rate so that the product kinetics resemble their data as closely as possible. Students are then asked to brainstorm all the ways this is a simplification of the real system (worksheet is provided as a Supplemental). This illustrates firsthand the subjectivity involved in attempting to produce a model of a system - both the biological concept model we began with, commonly shown in introductory courses, and the mathematical model developed in class.

### 3.1 Developing a model of catechol oxidase

A more complex model of catechol oxidase was constructed after the course was completed to analyze student data and determine the feasibility of parameter estimation with novice data. The simplified model of catechol oxidase (Eqn 5) was expanded using standard non-steady state mass-action kinetics. Catechol oxidase reversibly binds two substrates, catechol and oxygen, in random order addition. The presence of two substrates is something that students may not be actively considering, even though the enzymes function to catalyze an otherwise very slow reaction. This is likely due to the simplistic representation often shown in courses, such as Eqn. 1. The resulting system of equations has 11 parameters and 7 variables (Eqn 3.1).

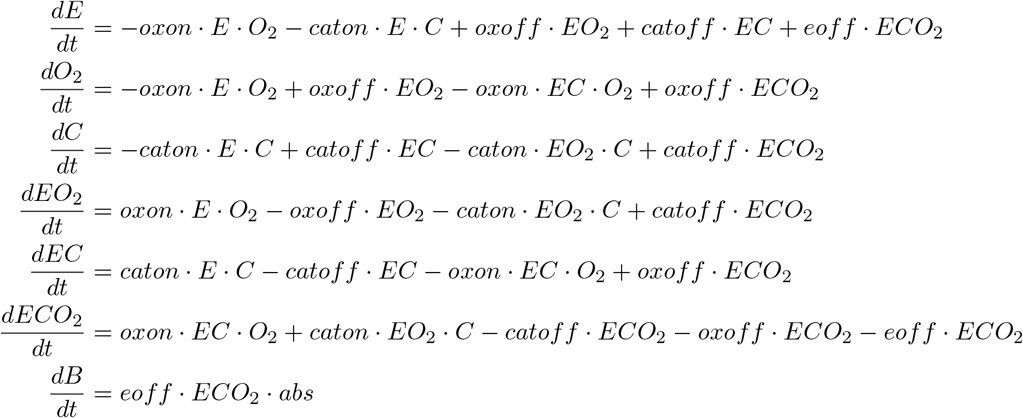

Here, oxygen (*O*_2_) binds (*oxon*) and unbinds (*oxoff*) the enzyme *E*. Catechol (*C*) binds (*caton*) and unbinds (*catoff*) the enzyme. Since this happens with random addition, we consider two intermediate species *EO*_2_ and *EC*. When both substrates are bound (*ECO*_2_) catalysis can occur (*eoff*), producing the product benzoquinone (*B*) which accumulates, darkening the reaction solution and enabling measurement via spectrophotometer. Concentration of *B* in terms of absorbance is approximated by the parameter *abs*. After catalysis, the enzyme is returned to the pool of free enzyme *E*.

Since this model was developed to handle novice student data, we included some additional parameters to control for common errors. Since the absolute concentration of enzyme or the substrates catechol and oxygen were not measured, we included volume to unit conversion factors. The amount of enzyme in a mililiter of potato slurry is *scaleE*. The amount of catechol substrate in a militer of catechol solution is *scaleC*. These factors may be somewhat variable across student groups, as pipetting accuracy varies, and the enzyme slurry is extracted from potatoes and contains some substrates that can oxidize. Students may fail to measure absorbance immediately after combining enzyme and substrate into the reaction mixture. To account for such delays, we included the parameter *shift*. Finally, we included a set of *scale* parameters to understand if pooling student data would reasonably agree with individual fits. The lower and upper bounds for the parameters were chosen based on literature when available. Parameter estimates are provided in Supplemental Table 4. We found reasonable agreement across parameter estimates, indicating the suitability of this approach for modeling and parameter estimation with student data (Fig. 2). The effect of varying enzyme concentrations is shown in Fig. 1, and the effect of varying catechol and oxygen concentrations are provided as Supplemental Fig. 5 and 6.

**Figure 1.**
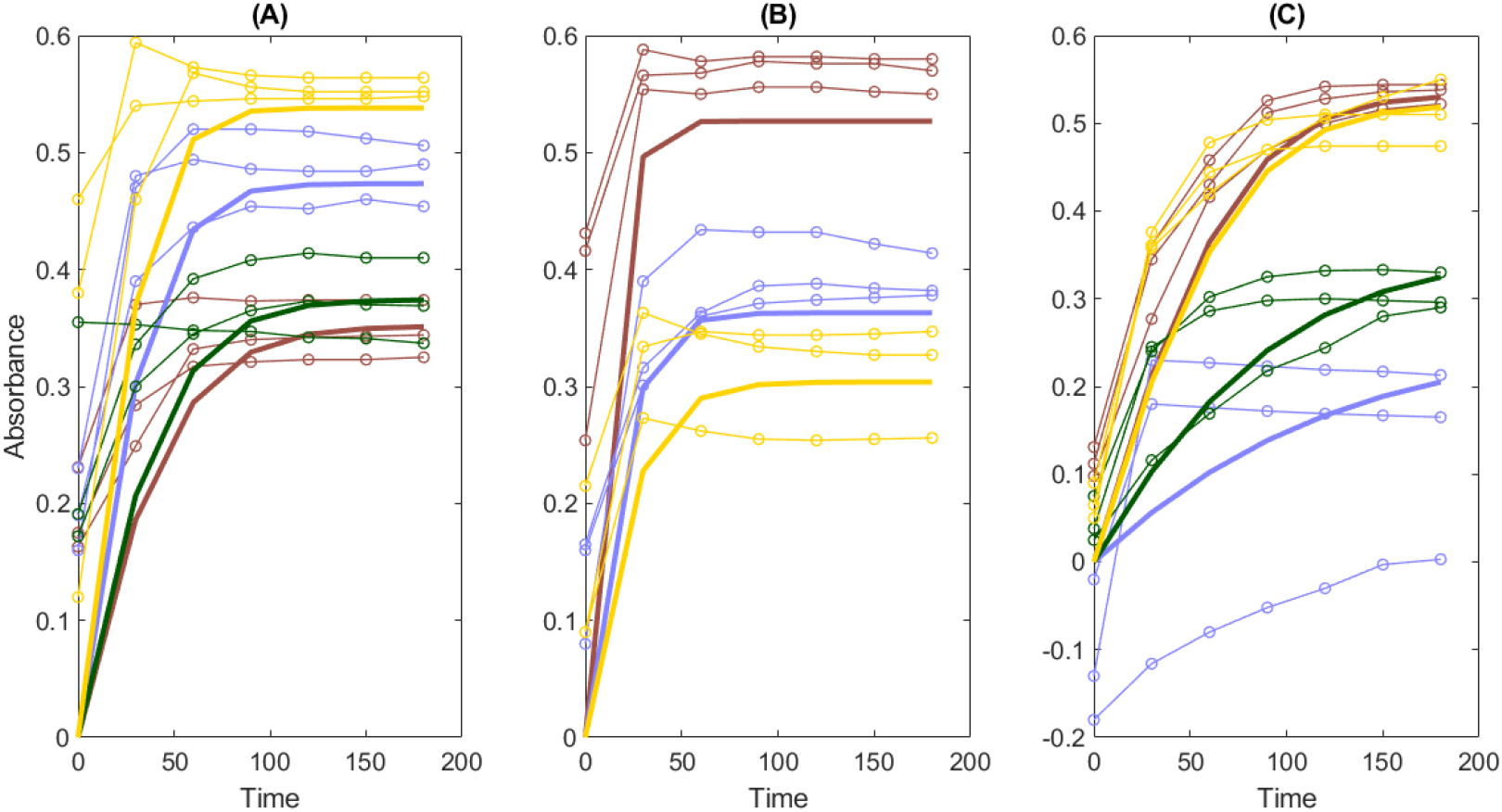
The complex mathematical model of catechol oxidase agrees with student data after parameter estimation. The mathematical model of catechol oxidase kinetics (bold) was fit to the data gathered across 4 groups of novice students (circles) to estimate kinetic parameters. Each color represents a different group. Students measured the kinetics of 1 mL (A), 2 mL (B), and 0.5 mL (C) of potato catechol oxidase slurry, modifying enzyme (catechol oxidase) concentration while holding substrate (catechol) constant.

**Figure 2.**
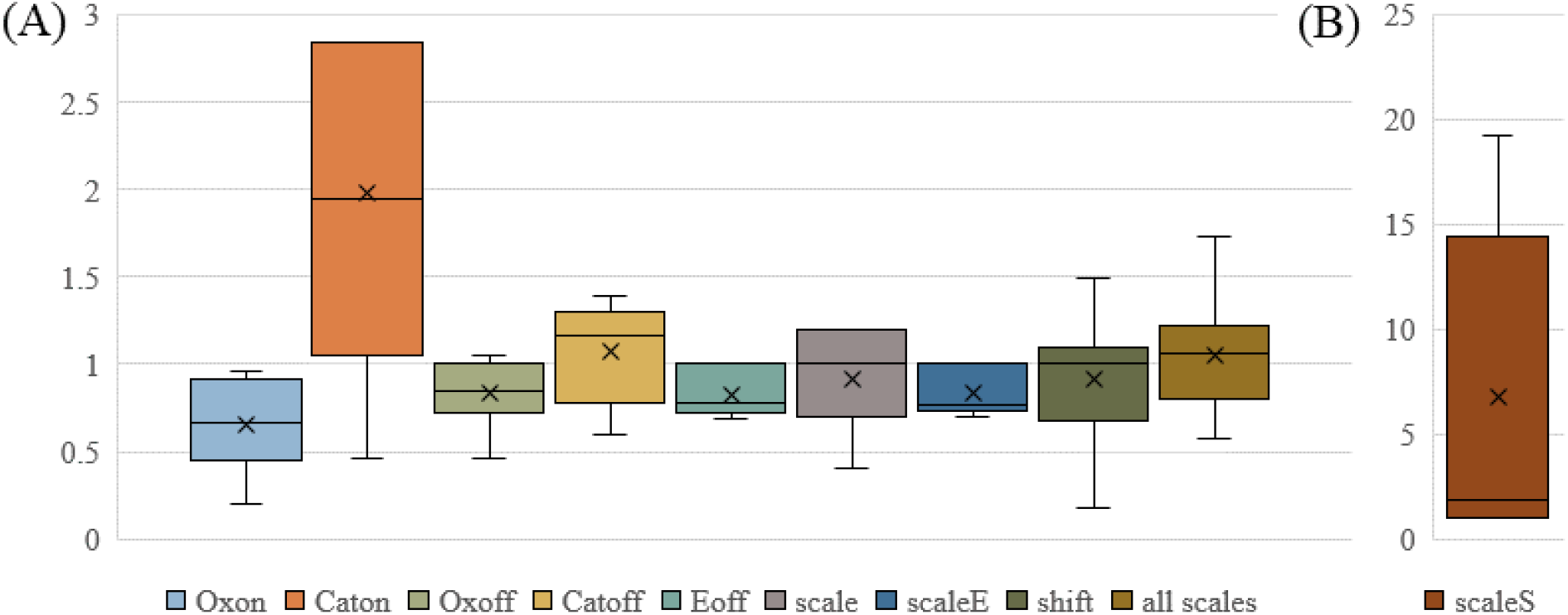
There was reasonable agreement between parameter estimates using data pooled across experiments and estimates from individual fits. Here we show the proportional difference, dividing individual fitted parameter values by the pooled parameter estimate. The biggest difference is seen in the *scaleS* parameter, which we use to approximate the concentration of substrate catechol. The parameter *Caton*, representing the binding of catechol, shows a wide spread but the values are small, ranging from 0.008 to 0.05.

### 3.2 Survey Results

Students were provided with surveys intended to track their opinions of math and biology before and after course completion. Survey items were selected for their relevance from existing instruments to capture opinion on math and biology [16, 17]. The range of possible responses was 1 (disagree) to 5 (agree). Three questions were significant with p*<*0.05 using Wilcoxon signed rank test, which is appropriate for ordinal data.

Among the math questions, question 1 *There is usually only one correct approach to solving a math problem*. was significant at the p*<*0.05 level (pre-test mean 2.03, post-test 1.64, indicating greater disagreement). An additional question (7) was significant at the p*<*0.1 level *Learning math changes my ideas about how the world works*. (pre-test mean 2.97, post-test mean 3.24, indicating greater agreement).

Among the biology questions, questions 13 *Reasoning skills used to understand biology can be helpful to my everyday life*. (Pre-test mean 4, post-test 4.30, indicating greater agreement) and 20 *I think about the biology I experience in everyday life*. (pre-test mean 3.12, post-test mean 3.52, indicating greater agreement) were both significant at the p*<*0.05 level. An additional question (15) was significant at the p*<*0.06 level *When studying biology, I relate the important information to what I already know rather than just memorizing it the way it is presented*. (pre-test mean 4.10, post-test mean 3.82, indicating greater disagreement).

At the end of the semester, students were asked to take an open-ended survey of their opinions on the course. Students enjoyed the experiments the most (29/35). Students disliked programming (12/35) and course assignments (11/35) the most. Students felt the course activities would help them understand math, biology, or programming better in the future (29/35), particularly programming (5/29). Most students felt the course was appropriately challenging (22/35). When asked if they would be interested to learn more about course topics, students mentioned biology (9/29, math (1/29), programming (5/29), and statistics (3/29).

## 4 Discussion

The tools discussed in this paper were implemented in a CURE introductory biology lab for freshmen students. During the course, students engaged in the modeling process, ranging from black box simulations of a simple model to model derivations, parameter estimation, and critique. The students also learned the experimental techniques involved in the catechol oxidase assay and some basic programming in R. Our findings indicate the feasibility of integrating modeling into existing cookbook labs, as well as the suitability of such activities for students with modeling-poor backgrounds.

Although these students had no prior knowledge of the course’s quantitative focus, and had little or no prior experience in modeling (e.g., differential equations, calculus, coding), we found that the presence of quant methods did not deter them. The majority of students found the course material appropriately challenging. Some students expressed interest to learn more math, programming, and statistics. Indeed, one such student worked on developing the complex model, parameter estimation, and this paper. The pre- and post-course surveys indicate that some of students’ opinions on math and biology changed as a result of completing the course. We found more significant changes in the biology survey questions, which was unexpected and may indicate the ability of math biology for students to appreciate biology. The lack of a control group limits the interpretability of these results. We weren’t able to determine what about the class might have caused these changes, but it would be interesting to pursue further in future studies. It seems likely that the emphasis on the variety of solutions to modeling a system, how models are inherently simplficiations, and critiquing the model may contribute positiviely to reduce math anxiety and increase ownership [9]. We hope to provide a starting point for future work on developing similar modules, including issues that we experienced during our work.

Finally, we were able to reasonably estimate parameters from student data acquired during the course. The parameter estimates were reasonably consistent, whether estimated by group or as a whole. This indicates the feasibility of integrating mathematical modeling into student experimental labs, including often challenging parameter estimation.

In conclusion, integrating mathematical modeling with student-driven experimentation is a feasible and effective way to bring quantitative skills to the classroom. The amount of time investment needed to accomplish this is typically a concern [5]. The original cookbook lab was 2 weeks, while the CURE lab was over a semester (approximately 10 weeks on material presented here). The tools presented here form a scaffolding that may be adapted to available time (Fig. 3), and similar approaches may be successful for other experimental systems and quantitative techniques.

**Figure 3.**
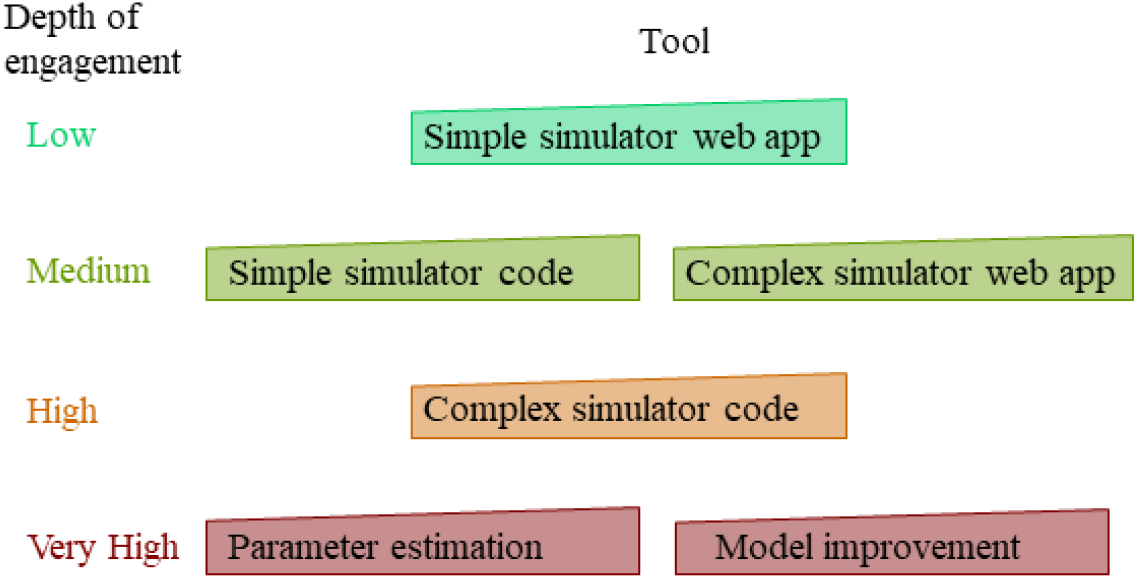
Different tools may be used according to time available and course objectives.

## Supporting information

Worksheet1

Worksheet2

Likert_Survey

Open_Survey

## ACKNOWLEDGMENTS

The authors would like to thank Dr Bill Wischusen and Dr Naohiro Kato for their suggestions and support during the development of the research question and course content. They also thank the students who provided valuable feedback, took the surveys, and tested the tools.

