## Supplemental for "Integrating Math Modeling, Coding, and Biology in a CURE Lab"

### 1 Data availability

Data on Likert survey responses, open survey responses, and the experimental data are available at 10.6084/m9.figshare.6169523. All data have been anonymized.

### 2 Likert survey on math and biology attitudes

| No. | Question | Mean pre-response | Mean post-response |
|---|---|---|---|
| 1 | There is usually only one correct approach to solving a math problem. | 2.03 | 1.64 |
| 2 | I do not expect formulas to help my understanding of mathematical ideas, they are just for doing calculations. | 2.12 | 1.82 |
| 3 | Math ability is something about a person that cannot be changed very much. | 2.12 | 2.15 |
| 4 | Nearly everyone is capable of understanding math if they work at it. | 4.06 | 3.88 |
| 5 | Understanding math means being able to recall something you've read or been shown. | 3.12 | 3.06 |
| 6 | In math, it is important for me to make sense out of formulas and procedures before I use them. | 4.12 | 4 |
| 7 | Learning math changes my ideas about how the world works. | 2.97 | 3.24 |
| 8 | Reasoning skills used to understand math can be helpful to me in my everyday life. | 3.55 | 3.58 |
| 9 | It is a waste of time to understand where math formulas come from. | 2.42 | 2.42 |
| 10 | Being good at math requires natural (i.e. innate, inborn) intelligence in math. | 2.3 | 2.45 |
| 11 | We use this statement to discard the survey of people who are not reading the questions. Please select Agree (not Strongly Agree) for this question. | 4.12 | 4.06 |
| 12 | To learn biology, I only need to memorize facts and definitions. | 2.08 | 2 |
| 13 | Reasoning skills used to understand biology can be helpful to my everyday life. | 4 | 4.30 |
| 14 | The subject of biology has little relation to what I experience in the real world. | 2 | 1.82 |
| 15 | When studying biology, I relate the important information to what I already know rather than just memorizing it the way it is presented. | 4.09 | 3.82 |
| 16 | There is usually only one correct approach to solving a biology problem. | 2.55 | 2.42 |
| 17 | Mathematical skills are important for understanding biology. | 3.39 | 3.42 |
| 18 | Biological principles are just to be memorized. | 2.03 | 2.09 |
| 19 | For me, biology is primarily about learning known facts as opposed to investigating the unknown. | 2.45 | 2.39 |
| 20 | I think about the biology I experience in everyday life. | 3.12 | 3.52 |
| 21 | I do not expect the rules of biological principles to help my understanding of the ideas. | 1.79 | 1.79 |

**Table 1.** A survey on opinions of math and biology were provided to the students before and after taking the course. A response of 1 indicates disagree and 5 agree. Questions 1 to 10 were taken from [1], and questions 11 to 21 from [2]. Note, question 11 has a typo (mentioning strongly agree), and students were verbally instructed to put 4.

### 2.1 Checking symmetry of the data

The survey data are 5-point Likert scale. Each respondent took pre- and post-tests. We checked the symmetry of the differences using histograms (Fig. 1). Overall, the data look symmetric.

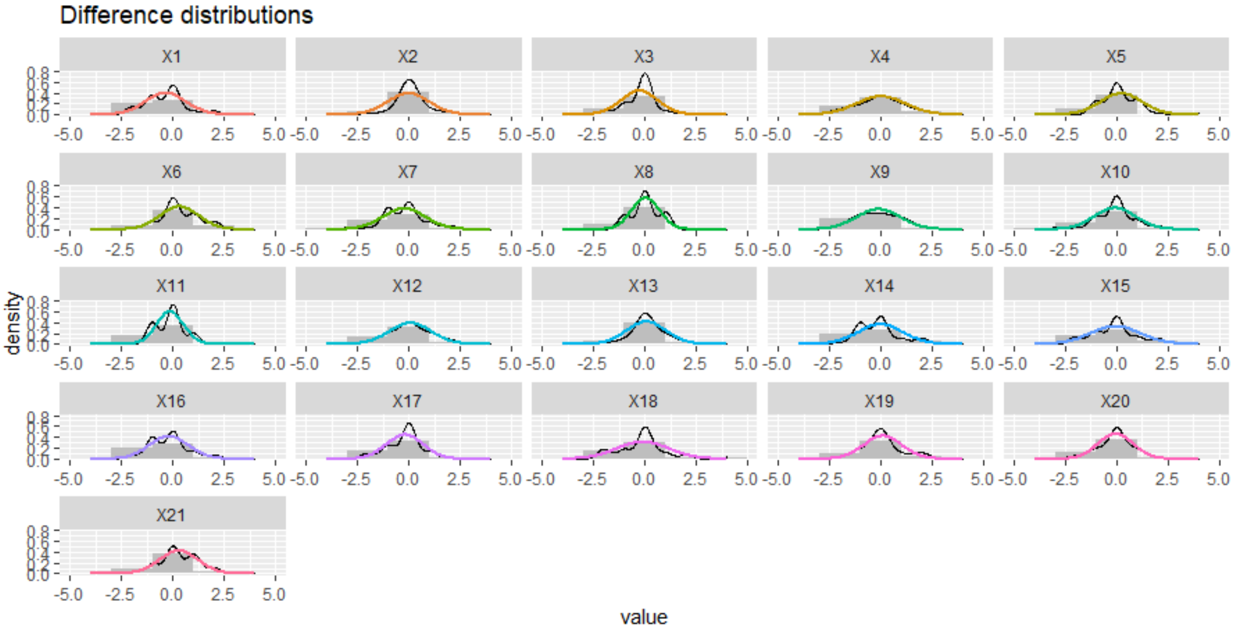

**Figure 1.** Distributions of the difference between the pre- and post-tests across the respondents. Distributions overall look symmetric.

### 2.2 Wilcoxon test results

Since the data are linked and ordinal, we used the non-parameteric Wilcoxon signed rank test in R 3.3.6 with the *wilcox_test* function. Results are in Table 2. Questions 1, 13, and 20 were significant at the $p<0.05$ level. The pre- and post-test response histograms for these three questions are provided in Fig. 2, 3, and 4.

| Q | group1 | group2 | n1 | n2 | statistic | p | p.signif |
|---|---|---|---|---|---|---|---|
| 1 | post | pre | 33 | 33 | 23.5 | 0.0339 | |
| 2 | post | pre | 33 | 33 | 63 | 0.195 | ns |
| 3 | post | pre | 33 | 33 | 107 | 0.952 | ns |
| 4 | post | pre | 33 | 33 | 39 | 0.215 | ns |
| 5 | post | pre | 33 | 33 | 141.5 | 0.812 | ns |
| 6 | post | pre | 33 | 33 | 81 | 0.568 | ns |
| 7 | post | pre | 33 | 33 | 111 | 0.0889 | ns |
| 8 | post | pre | 33 | 33 | 165.5 | 0.944 | ns |
| 9 | post | pre | 33 | 33 | 88.5 | 0.338 | ns |
| 10 | post | pre | 33 | 33 | 66 | 0.398 | ns |
| 11 | post | pre | 33 | 33 | 2.5 | 0.424 | ns |
| 12 | post | pre | 33 | 33 | 61.5 | 0.749 | ns |
| 13 | post | pre | 33 | 33 | 142.5 | 0.0349 | |
| 14 | post | pre | 33 | 33 | 43 | 0.182 | ns |
| 15 | post | pre | 33 | 33 | 28 | 0.0527 | ns |
| 16 | post | pre | 33 | 33 | 67 | 0.14 | ns |
| 17 | post | pre | 33 | 33 | 64 | 0.821 | ns |
| 18 | post | pre | 33 | 33 | 44 | 0.705 | ns |
| 19 | post | pre | 33 | 33 | 62.5 | 0.789 | ns |
| 20 | post | pre | 33 | 33 | 119.5 | 0.0383 | |
| 21 | post | pre | 33 | 33 | 33 | 1 | ns |

**Table 2.** Results of Wilcoxon test for pre- and post-test scores on Likert-scale questionnaire.

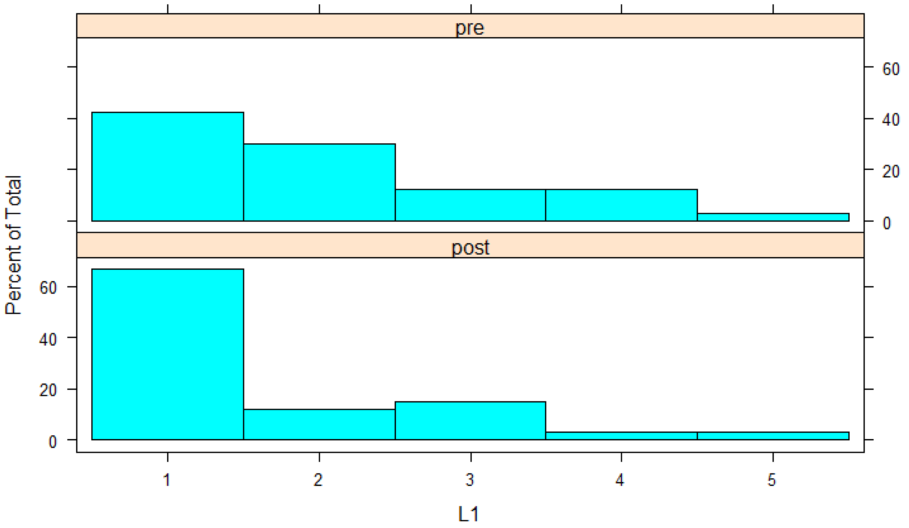

**Figure 2.** Question 1 pre- and post-test response distribution.

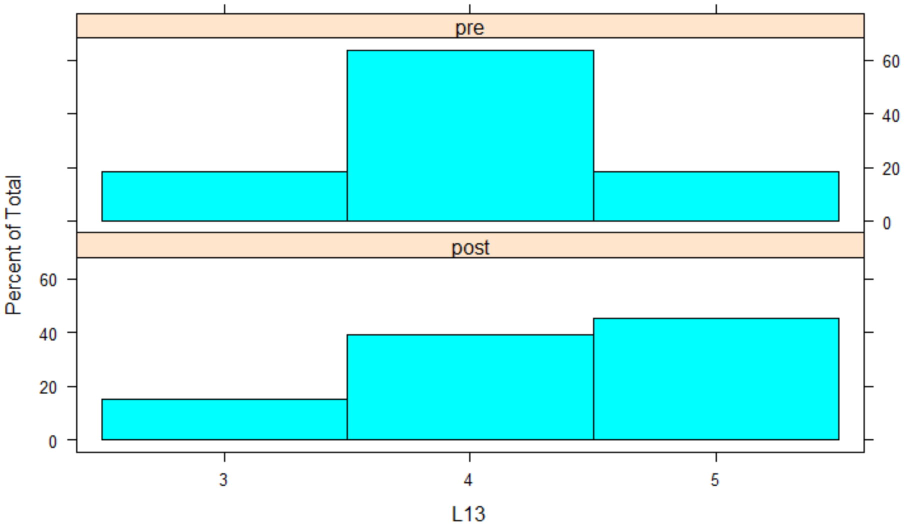

**Figure 3.** Question 13 pre- and post-test response distribution.

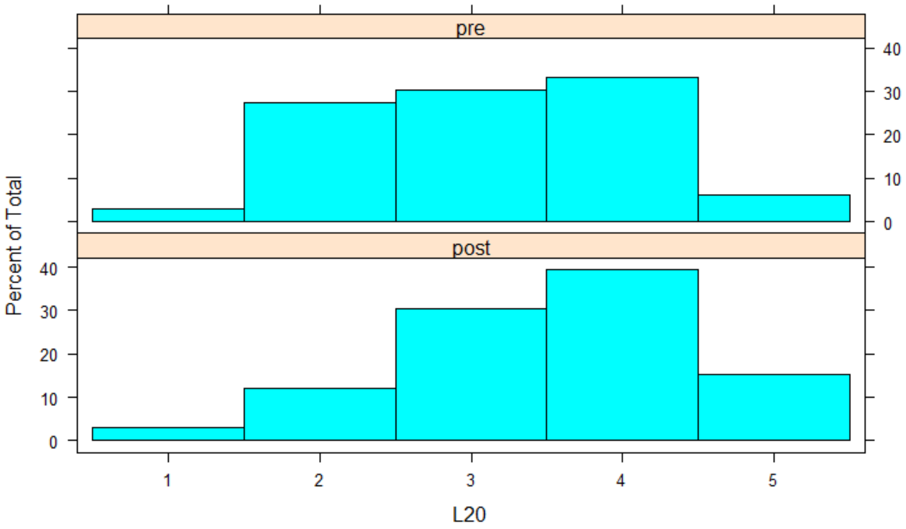

**Figure 4.** Question 20 pre- and post-test response distribution.

| | R | Adjusted |
|---|---|---|
| Pre-test | 0.37 | 0.54 |
| Post-test | 0.45 | 0.62 |
| All | 0.37 | 0.54 |

**Table 3.** Chronbach's Alpha results for the survey items indicate poor (<0.6) and acceptable (> 0.6) reliability. Adjusted using Spearman-brown.

#### 2.3 Checking survey items reliability

To calculate reliability, we used Chronbach Alpha. Questions 1-3,5,9,10,12,14,16,18,19 were reversed (where e.g., disagree = more math anxiety). The results are shown in Table 3.

| Parameter | Description |
| --- | --- |
| oxon | oxygen binding rate |
| caton | catechol binding rate |
| oxoff | oxygen unbinding rate |
| catoff | catechol unbinding rate |
| eoff | catalysis rate |
| abs | unit to absorbance conversion |
| scaleE | conversion, mL to units |
| scaleS | conversion, mL to units, *1000 |
| scaleO | conversion, mL to units (fixed at 1000) |
| shift | delay in beginning measurement, *30 seconds |
| scales | overall error in initial conditions (i.e., pipetting) |

**Table 4.** Parameters in the model were fit to student data. There was one *scales* parameter per each of the 25 student groups, to account for pipetting errors.

### 3 Complex Mathematical Model equations

$$
\begin{aligned}
\frac{dE}{dt} &= -Oxon \cdot E \cdot O_2 - Caton \cdot E \cdot C + Oxoff \cdot EO_2 + Catoff \cdot EC + Eoff \cdot ECO_2 \\
\frac{dO_2}{dt} &= -Oxon \cdot E \cdot O_2 + Oxoff \cdot EO_2 - Oxon \cdot EC \cdot O_2 + Oxoff \cdot ECO_2 \\
\frac{dC}{dt} &= -Caton \cdot E \cdot C + Catoff \cdot EC - Caton \cdot EO_2 \cdot C + Catoff \cdot ECO_2 \\
\frac{dEO_2}{dt} &= Oxon \cdot E \cdot O_2 - Oxoff \cdot EO_2 - Caton \cdot EO_2 \cdot C + Catoff \cdot ECO_2 \\
\frac{dEC}{dt} &= Caton \cdot E \cdot C - Catoff \cdot EC - Oxon \cdot EC \cdot O_2 + Oxoff \cdot ECO_2 \\
\frac{dECO_2}{dt} &= Oxon \cdot EC \cdot O_2 + Caton \cdot EO_2 \cdot C - Catoff \cdot ECO_2 - Oxoff \cdot ECO_2 - Eoff \cdot ECO_2 \\
\frac{dB}{dt} &= Eoff \cdot ECO_2
\end{aligned}
$$

### 4 Parameter Estimates

Approach to estimate parameters is described in main text. Parameter estimates by pooled and group fits is shown in Table 4.1, and the *scale* parameter estimates are shown in Tables 6 and 7.

#### 4.1 Fitting pooled and individual group data

When fitting pooled data, we do not apply individual error parameter *scales* to each group's data. Scales is applied to the initial conditions vector to account for experimental setup/pipetting errors. Instead, we use a single *scales* parameter. Model agreement to changes in concentration of catechol oxidase is shown in the main text, Fig. 1; agreement to changes in oxygen (Fig. 5) and catechol (Fig. 6) shown below.

| | Oxon | Caton | Oxoff | Catoff | Eoff | scale | scaleE | scaleS | shift |
|---|---|---|---|---|---|---|---|---|---|
| all pooled | 0.05 | 0.0176 | 12.0214 | 17.4819 | 6.9833 | 0.0005 | 9.9792 | 0.3 | 1 |
| catechol1 | 0.0458 | 0.05 | 10.1423 | 20.0816 | 5.1473 | 0.0005 | 8.0918 | 0.4164 | 1 |
| enzyme1 | 0.0224 | 0.0336 | 12.0776 | 23.4323 | 5.4083 | 0.0006 | 6.9219 | 5.781 | 0.3585 |
| enzyme2 | 0.0312 | 0.0081 | 10.2479 | 20.3551 | 4.7624 | 0.0004 | 7.2364 | 0.5679 | 1 |
| enzyme3 | 0.0225 | 0.0289 | 12.5799 | 24.2828 | 5.3058 | 0.0004 | 7.3649 | 3.5179 | 1.4889 |
| enzyme4 | 0.0331 | 0.0082 | 12.125 | 22.125 | 5.3971 | 0.0005 | 7.4986 | 1.9953 | 1.1884 |
| oxygen1 | 0.0482 | 0.05 | 8.2284 | 11.4627 | 7 | 0.0003 | 9.9994 | 0.3 | 1 |
| oxygen2 | 0.0098 | 0.05 | 5.538 | 10.3455 | 7 | 0.0006 | 9.9997 | 0.3 | 1 |
| oxygen3 | 0.0458 | 0.0343 | 10.1539 | 20.4462 | 4.9021 | 0.0002 | 7.6394 | 0.3 | 1 |
| oxygen4 | 0.0348 | 0.05 | 9.2326 | 15.6767 | 6.9766 | 0.0006 | 9.9915 | 5.1232 | 0.1725 |

**Table 5.** Parameter estimates obtained from fitting all data (*all pooled*) and fitting each group's data sets.

| | scale 1 | 2 | 3 | 4 | 5 | 6 | 7 | 8 | 9 | 10 |
|---|---|---|---|---|---|---|---|---|---|---|
| all pooled | 0.8697 | 0.8801 | 1.2302 | 0.7035 | 1.0538 | 1.0656 | 0.9469 | 0.726 | 0.5 | 0.7488 |
| catechol1 | 0.9731 | 1.0081 | 1.0365 | | | | | | | |
| enzyme1 | | | | 0.627 | 0.9368 | 0.9639 | | | | |
| enzyme2 | | | | | | | 1.3808 | 1.0216 | 0.622 | |
| enzyme3 | | | | | | | | | | 0.9249 |
| | 11 | 12 | 13 | 14 | 15 | 16 | 17 | 18 | 19 | 20 |
| all pooled | 0.6078 | 0.6945 | 1.0764 | 0.6078 | 1.0437 | 0.8786 | 0.5001 | 0.8268 | 0.5 | 0.5 |
| catechol1 | | | | | | | | | | |
| enzyme1 | | | | | | | | | | |
| enzyme2 | | | | | | | | | | |
| enzyme3 | 0.7241 | 0.8208 | | | | | | | | |
| enzyme4 | | | 1.1463 | 0.6207 | 1.0906 | | | | | |
| oxygen1 | | | | | | 1.5 | 0.6919 | 1.4424 | | |
| oxygen2 | | | | | | | | | 0.9507 | 0.6936 |
| | 21 | 22 | 23 | 24 | 25 | | | | | |
| all pooled | 0.5 | 1.5 | 1.1088 | 0.6823 | 1.5 | | | | | |
| catechol1 | | | | | | | | | | |
| enzyme1 | | | | | | | | | | |
| enzyme2 | | | | | | | | | | |
| enzyme3 | | | | | | | | | | |
| enzyme4 | | | | | | | | | | |
| oxygen1 | | | | | | | | | | |
| oxygen2 | 1.5 | | | | | | | | | |
| oxygen3 | | 0.8005 | 0.5791 | 1.5 | | | | | | |
| oxygen4 | | | | | 0.9901 | | | | | |

**Table 6.** Scale parameter estimates were fit for each group when fitting separately.

| | scale 1 | 2 | 3 | 4 | 5 | 6 | 7 | 8 | 9 | 10 |
|---|---|---|---|---|---|---|---|---|---|---|
| all pooled | 0.8697 | 0.8801 | 1.2302 | 0.7035 | 1.0538 | 1.0656 | 0.9469 | 0.726 | 0.5 | 0.7488 |
| catechol1 | 0.9731 | 1.0081 | 1.0365 | | | | | | | |
| enzyme1 | | | | 0.627 | 0.9368 | 0.9639 | | | | |
| enzyme2 | | | | | | | 1.3808 | 1.0216 | 0.622 | |
| enzyme3 | | | | | | | | | | 0.9249 |
| | 11 | 12 | 13 | 14 | 15 | 16 | 17 | 18 | 19 | 20 |
| all pooled | 0.6078 | 0.6945 | 1.0764 | 0.6078 | 1.0437 | 0.8786 | 0.5001 | 0.8268 | 0.5 | 0.5 |
| enzyme3 | 0.7241 | 0.8208 | | | | | | | | |
| enzyme4 | | | 1.1463 | 0.6207 | 1.0906 | | | | | |
| oxygen1 | | | | | | 1.5 | 0.6919 | 1.4424 | | |
| oxygen2 | | | | | | | | | 0.9507 | 0.6936 |
| | 21 | 22 | 23 | 24 | 25 | | | | | |
| all pooled | 0.5 | 1.5 | 1.1088 | 0.6823 | 1.5 | | | | | |
| oxygen2 | 1.5 | | | | | | | | | |
| oxygen3 | | 0.8005 | 0.5791 | 1.5 | | | | | | |
| oxygen4 | | | | | 0.9901 | | | | | |

**Table 7.** Scale parameter estimates were fit for each group when fitting separately.

#### 4.2 Model Fits to modifications of substrate (catechol and oxygen) concentrations

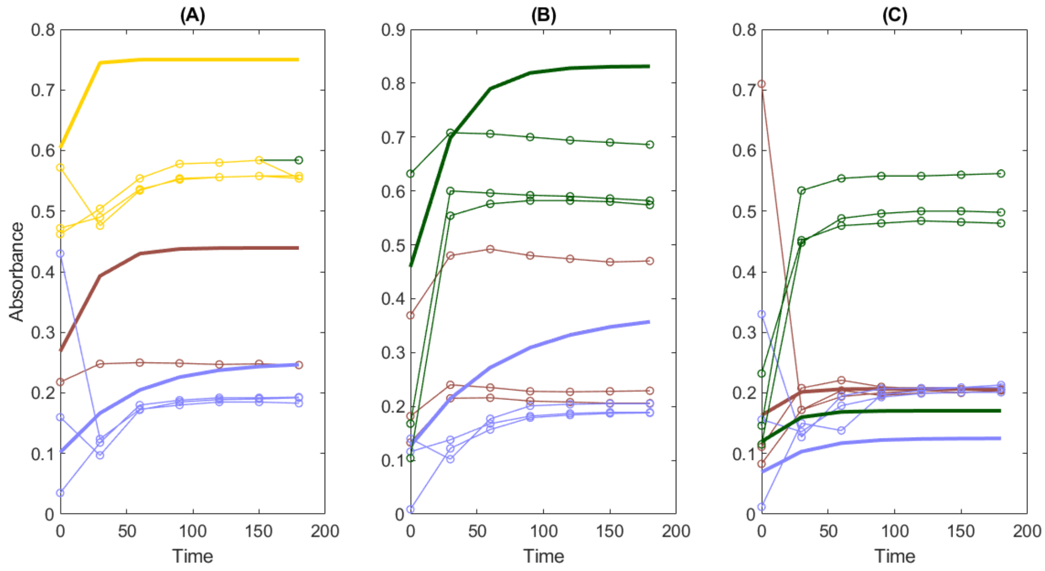

**Figure 5.** Effect of changing the oxygen concentration on the kinetics. Students measure dthe kinetics of dissolved oxygen (A), pipetting additional air into the solution (B), and covering the reaction tube (C). Each color represents a different group.

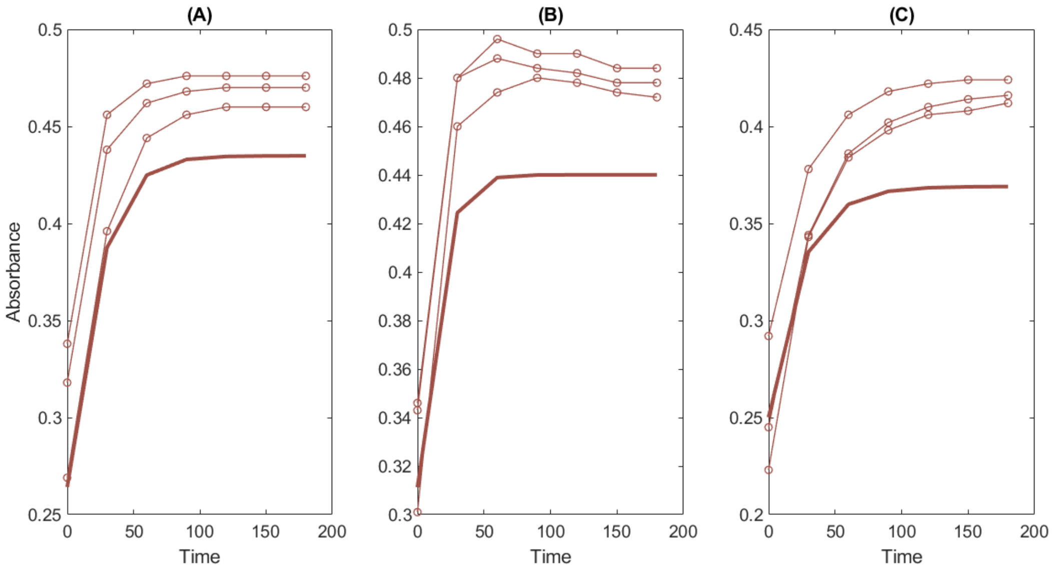

**Figure 6.** Effect of changing the catechol concentration on the kinetics. Students measured the kinetics of 5 mL (A), 7 mL (B), and 2 mL (C) of catechol. Each color represents a different group.

#### 4.3 Workflow to estimate parameters from student data

To begin, uncomment lines 50-56 in the curvefitting_code.m file. Comment line 59 to remove the fitted values presented in this paper. Matlab global optimization toolbox must be installed, or anotehr fitter should be used.
