## Supplementary material for "Integrating Math Modeling, Coding, and Biology in a CURE Lab": Worksheet1

1. Fill in the reaction rates for your experiments in the table below. Indicate which variable you altered in experiments 1 and 2.

| | Linear Regression | Final/initial | Plateau |
|---|---|---|---|
| Control | | | |
| Exp1 | | | |
| Exp2 | | | |

Was the slope of the linear regression significant? (Yes No)

Does the linear regression look like your data? (Yes No)

2. The 3 methods we used to get the reaction rate aren't perfect. Write one of the problems next to the methods below:

Linear Regression ___________________________________________________________________________

Final/Initial ________________________________________________________________________________

Plateau _____________________________________________________________________________________

3. Run the simulation and compare the results to your data. Does it look the same? (Yes No)

4. What kind of stuff do you think the model is simulating?

5. An enzyme reaction occurs when you add an enzyme with its two substrates. Then it can catalyze the reaction between the substrates to make the product.

What decreases in an enzyme reaction?

What increases?

**6. Rate** means how quickly something happens. An example of a rate is: __________________________

If you measured a rate, it would have these units:

7. Let's say our enzyme has a reaction rate of 1 per second. This the enzyme can create one molecule of product every second. How many substrates will I lose after 1 second if I add an enzyme?

How many product molecules will I gain in 1 second?

8. What if I have 2 enzymes? How many substrates will I lose in 1 second?

How many products will I gain in 1 second?

9. Let's say our rate is ***r*** per second. This means every second I will lose _____ substrates and gain ____ products.

10. The substrate is **decreasing** at a rate of ***r*** per second.

Write as an equation:

11. What will happen if we add 2 enzymes?

Or ***E*** enzymes?

Now our equation is:

12. We know the product must be increasing at the same rate. Write the equation for the product:

13. The substrate is **changing over time.** It is **decreasing** at the rate **r per second** for each **enzyme** molecule.

Mathematically:

Means

14. Now write the equations for the enzyme:

S

P

E

To complete the equations:

- Does the enzyme change over time?
- What happens when we run out of substrate?

ICA 1 biol 1208

The model is simplified. For each of the simplifications, write how it might be important. Did the scientific article you read mention any of these?

1. Oxygen

2. Temperature

3. pH

4. Enzyme stability

5. Inhibition by the product

6. _________________

7. _________________

Pick one with your lab partner to research for your experiment next week
