## Supplementary material for "Integrating Math Modeling, Coding, and Biology in a CURE Lab": Worksheet2

ICA 5. Coding & Modeling Worksheet Name:

Fill out the first 4 columns of the following table:

| Experiment | Catechol volume | Enzyme volume | O2 volume | F estimate | R estimate |
|---|---|---|---|---|---|

Fill in the comments on the code (indicated by #). Then modify the parameters “f” and “r” so that the simulation matches your data.

1. Why do we need the parameter “f”?

2. What does “f” control in the model?

3. Why do we need the parameter “r”?

4. What does “r” control in the model?

5. Can you find one “f” value and 6 “r” values so that the simulation matches your data? (Yes or no)

6. Do you think the model needs to be modified? If so, what should it include?

Modifying the model.

Run the simulations for inhibition and oxygen.

1. What effect does inhibition have on the simulation’s shape?

2. What effect does including oxygen have on the simulation’s shape?

3. What happens when you change the default parameter numbers? Can you make the effects disappear/have them make a larger difference?

4. Do either of these models seem more plausible or realistic?

5. How would you modify the model to incorporate Temperature?

6. How would you modify the model to incorporate pH?
