## Supplementary material for "Integrating Math Modeling, Coding, and Biology in a CURE Lab": Likert_Survey

Circle the number that best describes how well you agree with the statement.

| | Disagree | | | | Agree |
|---|---|---|---|---|---|
| 1. There is usually only one correct approach to solving a math problem. | 1 | 2 | 3 | 4 | 5 |
| 2. I do not expect formulas to help my understanding of mathematical ideas, they are just for doing calculations. | 1 | 2 | 3 | 4 | 5 |
| 3. Math ability is something about a person that cannot be changed very much. | 1 | 2 | 3 | 4 | 5 |
| 4. Nearly everyone is capable of understanding math if they work at it. | 1 | 2 | 3 | 4 | 5 |
| 5. Understanding math means being able to recall something you've read or been shown. | 1 | 2 | 3 | 4 | 5 |
| 6. In math, it is important for me to make sense out of formulas and procedures before I use them. | 1 | 2 | 3 | 4 | 5 |
| 7. Learning math changes my ideas about how the world works. | 1 | 2 | 3 | 4 | 5 |
| 8. Reasoning skills used to understand math can be helpful to me in my everyday life. | 1 | 2 | 3 | 4 | 5 |
| 9. It is a waste of time to understand where math formulas come from. | 1 | 2 | 3 | 4 | 5 |
| 10. Being good at math requires natural (i.e. innate, inborn) intelligence in math. | 1 | 2 | 3 | 4 | 5 |
| 11. We use this statement to discard the survey of people who are not reading the questions. Please select Agree (not Strongly Agree) for this question. | 1 | 2 | 3 | 4 | 5 |

| | Disagree | | | | Agree |
|---|---|---|---|---|---|
| 12. To learn biology, I only need to memorize facts and definitions. | 1 | 2 | 3 | 4 | 5 |
| 13. Reasoning skills used to understand biology can be helpful to my everyday life. | 1 | 2 | 3 | 4 | 5 |
| 14. The subject of biology has little relation to what I experience in the real world. | 1 | 2 | 3 | 4 | 5 |
| 15. When studying biology, I relate the important information to what I already know rather than just memorizing it the way it is presented. | 1 | 2 | 3 | 4 | 5 |
| 16. There is usually only one correct approach to solving a biology problem. | 1 | 2 | 3 | 4 | 5 |
| 17. Mathematical skills are important for understanding biology. | 1 | 2 | 3 | 4 | 5 |
| 18. Biological principles are just to be memorized. | 1 | 2 | 3 | 4 | 5 |
| 19. For me, biology is primarily about learning known facts as opposed to investigating the unknown. | 1 | 2 | 3 | 4 | 5 |
| 20. I think about the biology I experience in everyday life. | 1 | 2 | 3 | 4 | 5 |
| 21. I do not expect the rules of biological principles to help my understanding of the ideas. | 1 | 2 | 3 | 4 | 5 |
