## Supplementary material for "Integrating Math Modeling, Coding, and Biology in a CURE Lab": Open_Survey

### End of semester survey

1. What was your favorite thing we did this semester?
2. What was your least favorite thing we did this semester?
3. Do you think the activities we did in the course will help you understand math, biology, or programming better in the future?
4. Do you think the course was too hard or too easy? Why?
5. If you have any feedback for the TA, please enter it below.
6. Would you be interested in learning more about modeling, programming, or statistics?
